# Peripheral Myeloid-Derived Suppressor Cells are good biomarkers of the efficacy of Fingolimod in Multiple Sclerosis

**DOI:** 10.1101/2022.08.22.504792

**Authors:** Celia Camacho-Toledano, Isabel Machín-Díaz, Leticia Calahorra, María Cabañas, David Otaegui, Tamara Castillo-Triviño, Luisa María Villar, Lucienne Costa-Frossard, Manuel Comabella, Luciana Midaglia, José Manuel García-Domínguez, Jennifer García-Arocha, María Cristina Ortega, Diego Clemente

**Affiliations:** Neuroimmuno-Repair Group, Hospital Nacional de Parapléjicos-SESCAM, Finca La Peraleda s/n, 45071.Toledo, Spain; Multiple Sclerosis Unit, Biodonostia Health Institute, 20014. Donostia-San Sebastián, Spain; Neurology Department, Hospital Universitario Donostia, San Sebastián, Spain; Immunology Department, Hospital Universitario Ramón y Cajal, Instituto Ramón y Cajal de Investigación Sanitaria (IRYCIS), Madrid, Spain; Multiple Sclerosis Unit, Neurology, Ramón y Cajal University Hospital, Madrid, Spain; Neurology-Neuroimmunology Service, Centre d’Esclerosi Múltiple de Catalunya (Cemcat), Institut de Recerca Vall d’Hebron, Hospital Universitari Vall d’Hebron, Universitat Autònoma de Barcelona, Barcelona, Spain; Multiple Sclerosis Unit. Department of Neurology, Hospital General Universitario Gregorio Marañón, Madrid, Spain

## Abstract

The increasing number of treatments that are now available to manage patients with multiple sclerosis (MS) highlights the need to develop biomarkers that can be used within the framework of individualized medicine. Fingolimod is a disease-modifying treatment that belongs to the sphingosine-1-phosphate receptor modulators. In addition of inhibiting T cell egression from lymph nodes, fingolimod promotes the immunosuppressive activity of Myeloid-Derived Suppressor Cells (MDSCs), a cell type that can be used as a biomarker of disease severity, and of the degree of demyelination and extent of axonal damage in MS. In the present study, we have assessed whether the abundance of circulating monocytic-MDSCs (M-MDSCs) may represent a useful biomarker of fingolimod efficacy. As such, blood immune cells were analyzed at disease onset in the experimental autoimmune encephalomyelitis (EAE) MS mouse model. Fingolimod treated animals presented a milder EAE course with less demyelination and axonal damage, although a few animals did not respond well to treatment and they invariably had fewer M-MDSCs prior to initiating the treatment. Remarkably, M-MDSC abundance was also found to be an important and specific parameter to distinguish EAE mice prone to better fingolimod efficacy. Finally, in a translational effort, M-MDSCs were quantified in MS patients at baseline and correlated with different clinical parameters after 12 months of fingolimod treatment. The data obtained indicated that the M-MDSCs at baseline were highly representative of a good therapeutic response to fingolimod, i.e. patients who met at least two of the criteria used to define non-evidence of disease activity (NEDA-3) 12 months after treatment, providing relevant information of intention-to-treat MS patients. Collectively, our data indicate that M-MDSCs might be a useful predictive biomarker of the response of MS patients to fingolimod.

## 1. Introduction

Multiple sclerosis (MS) is a chronic inflammatory demyelinating disease of the central nervous system (CNS) that affects about 2.8 million people worldwide [1]. However, the development of MS does not follow a unique pattern but rather it has an unpredictable course. Around 85% of patients develop the relapsing-remitting (RR-MS) form of the disease, which is essentially characterized by episodes of neurological dysfunction (relapses) followed by partial or full recovery [2]. Our current understanding of the influence of the immune system on MS pathology has led to the approval of 17 disease modifying treatments (DMTs) for RR-MS, providing neurologists and patients a wide range of therapeutic options [3]. Clinicians must select treatment strategies for each individual based on a personalized risk and benefit assessment [4]. However, while the therapeutic options are constantly evolving, escalation is still a common strategy followed in the decision-making process for MS treatment [5]. It consists in using a first-line low risk, moderate efficacy therapy, and then monitoring the effectiveness of the treatment. When the DMT fails in controlling the disease activity, it is switched to a more efficacious one. This approach is quite conservative and it is most applicable to patients with a lower risk of developing an aggressive form of MS [6]. Mounting evidence indicates that early implementation of efficacious treatment may prevent chronic immune activation and better preserve long-term neural integrity [7]. Indeed, the most effective drugs should be indicated in patients with very active MS associated with an earlier development of disability. Among other DMTs, those designed to act as sphingosine-1-phosphate receptor (S1PR) modulators are a large family of drugs that has the ability to be used for this purpose.

Fingolimod (FTY720), known commercially as Gilenya^TM^, is a S1PR antagonist that was the first oral DMT approved for the treatment of RR-MS [8]. It is a structural analogue of sphingosine that upon phosphorylation acts as a modulator of S1PR1, 3, 4 and 5, with higher affinity for S1PR1. Fingolimod is a potent inhibitor of lymphocyte egression from lymphoid tissues as a result of the internalization and degradation of S1PR1 [9], although additional biological effects on other CNS or immune cells have also been described [10]. The heterogeneity of MS makes it difficult and costly to determine the effectiveness of any specific treatment, including fingolimod. Hence, it is important to identify biomarkers that provide information about disease progression and treatment efficacy for each individual to avoid undesirable side effects, and to reduce the cost burden to Health Systems. In this sense, a number of biomarkers have been described to monitor the effects of exposure to a drug and its pharmacodynamics, as well as to identify the likelihood of toxicity as an adverse effect of fingolimod treatment [11–13]. In recent years, the role of cells of the adaptive immune response as predictive biomarkers to anticipate the most positive response to fingolimod in MS patients has been addressed [14–16]. However, there is little data regarding the innate immune cells to predict the responsiveness of MS patients to fingolimod, attributed to CD56^bright^ NK cells [17], with no data regarding the myeloid cell compartment.

Myeloid-derived suppressor cells (MDSCs) are a heterogeneous population generated in the bone marrow during myelopoiesis that contribute to the innate immune response. Under pathological conditions, blocking their differentiation in conjunction with exposure to inflammatory mediators activates these cells, provoking their acquisition of immunosuppressive functions [18, 19]. In the experimental autoimmune encephalomyelitis (EAE) mouse model of MS, splenic MDSCs are composed of two subpopulations: polymorphonuclear-MDSCs (PMN-MDSCs, CD11b^+^Ly-6C^int^Ly-6G^+^) and monocytic-MDSCs (M-MDSCs, CD11b^+^Ly-6C^hi^Ly-6G^-^, which are phenotypically indistinguishable from Ly-6C^hi^ inflammatory monocytes in blood, i.e. Ly-6C^hi^ cells [20]). The abundance of M-MDSCs has repeatedly been related to disease severity in the EAE model [21–24]. Indeed, in several immune-mediated animal models other than EAE, it appears that fingolimod or other S1PR agonists may be implicated in the infiltration capacity, the expansion and the immunosuppressive activity of MDSCs through their antagonism of S1PR1, 3 and 5 [25–28]. Hence, there appears to be a clear relationship between the therapeutic response to fingolimod and the abundance of MDSCs. However, the utility of these cells as potential biomarkers of the response to fingolimod in inflammatory conditions (including EAE and MS) remains totally unexplored. Circulating PMN-MDSCs (identified as CD11b^+^CD33^+^HLA-DR^-/low^CD14^-^CD15^+^LOX1^+^ cells) and M-MDSCs (CD11b^+^CD33^+^HLA-DR^-/low^CD14^+^CD15^-^ cells) [29], yet MDSCs have been little studied in MS and there is conflicting data as to the cell type most strongly involved or altered during the clinical course of MS [30–32]. There is also little information on the possible effect of DMTs on MDSCs, most of which has been limited to glatiramer acetate (GA), which has been reported to induce a negligible increase in MDSCs [32]. Notably, there is no data about MDSCs as a predictive biomarker for MS treatments.

In the current work, we present novel data in the context of EAE and MS patients indicating that M-MDSCs could serve as a biomarker for responsiveness to fingolimod treatment and its efficacy in EAE. Patients who exhibited a good therapeutic response to fingolimod at 12 months of treatment had a greater level of circulating M-MDSCs at baseline relative to those considered to be non-responders. Our observations in EAE and MS might serve to identify new tools that in turn, could help modify the prescribing criteria of neurologists’ and ultimately, patients’ quality of life.

## 2. Materials and methods

### 2.1. Induction of EAE

Chronic progressive EAE was induced by subcutaneous immunization of female 6- to 8-week-old C57BL/6 mice (Janvier Labs, France) with 200 µL of a Myelin Oligodendrocyte Glycoprotein solution (200 µg,MOG_35-55_ peptide: Genscript, New Jersey, USA) emulsified in complete Freund’s Adjuvant (CFA) that contained 4 mg/mL of *Mycobacterium tuberculosis* (BD Biosciences, Franklin Lakes, New Jersey, USA). Immunized mice were administered Pertussis toxin intravenously (iv) through the tail vein (250 ng/mouse: Sigma Aldrich, St. Louis, MO, USA) in a final volume of 100 µL on the day of immunization and 48 hours later. EAE was scored clinically on a daily basis by three observers in a double-blind manner as follows: 0, no detectable signs of EAE; 0.5, half limp tail; 1, whole limp tail; 1.5, hind limb inhibition (unsteady gait and/or poor hind leg grip); 2, hind limb weakness; 2.5, bilateral partial hind limb paralysis or unilateral full hind limb paralysis; 3, complete bilateral hind limb paralysis (paraplegia); 3.5, partial forelimb paralysis; 4, tetraplegia; 4.5, moribund; and 5, death. In accordance with ethical regulations, humane end-point criteria were applied when an animal reached a clinical score ≥4, when clinical score ≥3 was reached for more than 48 h, or when self-mutilation was evident, persistent retention of urine, 35% weight loss and signs of stress or pain for more than 48 h, even if the EAE score was <3. The generation of sounds, lordokyphosis or stereotypic behavior were considered as signs of stress. In the present study, no animals presented signs of pain or stress, and none reached a clinical score >3.5.

The individualized treatment and follow-up was carried out as follows. The first day clinical signs of EAE were detected (“onset”, therapeutic window during which the clinical score debuts between 0.5 and 1.5), each animal was administered fingolimod orally (N = 18; 3 mg/Kg in distilled water: EAE-FTY720, kindly provided by Novartis Pharma AG, Switzerland [33–35]) or the vehicle alone (N =19; distilled water: EAE-Veh) on a daily basis for 14 days. All mice were sacrificed on the fifteenth day post-onset (dpo) and four different variables were analyzed in all the EAE mice: a) the clinical signs; b) the peripheral blood content; c) splenic content; and d) histopathological parameters.

All the animal manipulations were approved by the institutional ethical committee (*Comité Ético de Experimentación Animal del Hospital Nacional de Parapléjicos*), and all the experiments were performed in compliance with the European guidelines for animal research (EC Council Directive 2010/63/EU, 90/219/EEC, Regulation No. 1946/2003), and with the Spanish National and Regional Guidelines for Animal Experimentation (RD 53/2013 and 178/2004, Ley 32/2007 and 9/2003, Decreto 320/2010).

### 2.2. Evaluation of the clinical signs

The clinical signs were established through observational evaluation of the functional behavior of each mouse, allowing different parameters of the EAE clinical course to be defined: i) the day of onset, the first day mice had a clinical score ≥0.5; ii) clinical score at onset; iii) the maximum clinical score; iv) the day when animals reached the maximum clinical score or “peak” defined as the first day a clinical score ≥1.5 was repeated; v) the severity index, the maximum clinical score/the days elapsed from onset to peak [24]; vi) the accumulated clinical score from onset to peak and the total accumulated clinical score, considered as the sum of the individual clinical scores from the day of onset until the peak or the end of the treatment, respectively; vii) the percentage recovery [(the maximum clinical score – the residual score at the end of treatment) x 100/maximum clinical score]; viii) the recovery index [(the maximum clinical score at peak – the residual score at the end of treatment)/ days elapsed from the peak to the recovery]; and ix) the residual score, the clinical score at the end of the follow-up.

### 2.3. Flow cytometry analysis of peripheral blood and splenocytes

Blood was obtained from the retro-orbital venous sinus of isofluorane anesthetized mice at disease onset and collected in 2% EDTA tubes, and the erythrocytes were then lysed in a 15 mL tube with 2.5 mL ACK lysis buffer: 8.29 g/L NH_4_Cl; 1 g/L KHCO_3_; 1 mM EDTA in distilled H_2_O at pH 7.4 (Panreac). Subsequently, the lysis reaction was stopped by adding wash buffer (sterile 1X Phosphate Buffered Saline-PBS) and the blood cells were recovered by centrifugation at 210 g for 5 min at room temperature (RT). The Fc cell receptors were then blocked for 10 min at 4°C with an anti-CD16/CD32 antibody (10 μg/mL: BD Biosciences 553142) and the cells were resuspended in 25 μL of staining buffer: PBS supplemented with 10% heat-inactivated fetal bovine serum (FBS: Linus); 25 mM HEPES buffer (Gibco); and 2% Penicillin/Streptomycin (P/S: Gibco). The cells were further labelled for 20 min at 4°C in the dark with the following fluorochrome-conjugated monoclonal antibodies diluted in 25 μL of staining buffer (Suppl. Table 1): anti-Ly-6C-FITC (AL-21 clone), anti-Ly-6G-PE (1A8 clone), anti-CD11b-PerCP-Cy5.5 (M1/70 clone: all from BD Biosciences); anti-MHC-II-PE-Cy7 (M5/114.15.2 clone), anti-CD11c-APC (N418 clone), anti-F4/80-eFluor450 (BM8 clone) to stain myeloid cells (eBioscience-Thermo Fisher Scientific); and anti-CD3e-PB (500A2 clone), anti-CD4-PE (RM4-5 clone), anti-CD8a-FITC (53-6.7 clone: all from BD Bioscience), anti-CD25-BV421 (PC61.5 clone) and anti-CD69-APC (H1.2F3 clone) for lymphoid cell staining (Invitrogen). The blood cells were then rinsed with staining buffer, recovered by centrifugation at 210 g for 5 min, fixed for 10 min at RT with 4% paraformaldehyde (PFA: Sigma-Aldrich) in 0.1 M Phosphate Buffer (pH 7.4, PB), resuspended in PBS and finally analyzed in a FACS Canto II cytometer (BD Biosciences) at the Flow Cytometry Core Facility of the *Hospital Nacional de Parapléjicos*. The data obtained was analyzed using the FlowJo 10.6.2 software (Tree Star Inc.).

Fresh spleens were collected from euthanized mice at the end of fingolimod/vehicle administration and the tissue was processed mechanically to obtain a single cell suspension that was passed through a 40 µm filter (BD Biosciences), before it was washed in a 50 mL tube with cold supplemented RPMI medium: RPMI 1640 (Gibco) with 2mM L-Glutamine (L-Glu: Thermofisher), 10% FBS and 1% P/S. Erythrocytes were lysed with 1 mL of ACK lysis buffer and the reaction was stopped by adding PBS. After that, splenocytes were recovered by centrifugation at 210 g for 5 min and immunostained with myeloid/lymphoid markers following the same protocol as for the peripheral blood samples.

### 2.4. Tissue extraction and histopathological analysis

After removing their spleen, all the animals were perfused transcardially with 2% PFA, and their spinal cords were dissected out and post-fixed for 4 h at RT in the same fixative. After immersion in 30% (w/v) sucrose in PB for 12 h, coronal cryostat sections (20 µm thick: Leica, Nussloch) were thaw-mounted onto Superfrost® Plus slides. A randomly selected group of 18 EAE mice from the main cohort previously used for the spleen analysis was studied, obtaining histopathological parameters from the spinal cord: the area and percentage of demyelinated white matter, and the axonal damage associated to white matter lesions.

Demyelinated regions were quantified in eriochrome cyanine (EC) stained spinal cord tissue. Sections of the spinal cord were air-dried at RT for 12 h followed by 2 h at 37°C, placed in fresh acetone for 5 min and air-dried again for 30 min before they were stained for 30 min with 0.2% EC solution. The sections were then differentiated in 5% alum and borax-ferricyanide for 10 min each, and after rinsing with abundant tap water, correct differentiation was assessed under the microscope to ensure myelinated areas were stained blue and demyelinated areas appeared yellowish-white. Finally, the slides were dehydrated in a graded ethanol series and xylol solutions, and they were mounted with Entellan (Merck) for preservation.

To analyze axonal damage, the non-phosphorylated form of the neurofilament protein (SMI-32) was stained by immunohistochemistry. After several rinses in PB, the sections were treated with 10% methanol in PB for 20 min and then incubated for 1 h at RT in incubation buffer: 5% normal donkey serum (NDS: Merck) and 0.2% Triton X-100 (Merck) diluted in PBS. Immunostaining was performed by incubating the sections overnight at 4°C with the SMI-32 primary antibody (Biolegend: Suppl. Table 1) in incubation buffer. The sections were then incubated for 1 h at RT with the corresponding fluorescent secondary antibody diluted in staining buffer (1:1000: Invitrogen), the cell nuclei were stained with Hoechst (10 µg/mL: Sigma Aldrich) and the samples were mounted in Fluoromont-G (Southern Biotech). Omission of the primary antibody as a control staining yielded no signal.

### 2.5. Image acquisition and analysis

To quantify the degree of demyelination, sections were selected from the randomly subcohort group of 18 EAE mice (at least 3 sections per mouse, each separated by 340 µm). EC stained spinal cord sections were analyzed on a stereological Olympus BX61 microscope to quantify demyelination, using a DP71 camera (Olympus) and VisionPharm software for anatomical mapping. Superimages were acquired at a 10X magnification using the mosaic tool and analyzed with ImageJ, expressing the results as the total area of white matter demyelination and relative to the whole white matter area.

To quantify axonal damage, 20X tile scan images from the spinal cord of each animal were obtained on an IX83 inverted microscope (Olympus) equipped with CellSense Dimensions software for large-area imaging. The area of axon damage relative to the whole spinal cord or the total demyelinated area was analyzed with an *ad-hoc* plug-in designed by the Microscopy and Image Analysis Core Facility at the *Hospital Nacional de Parapléjicos*. After selecting the demyelinated area, a threshold for immunofluorescence was established and SMI-32 immunostaining was measured, representing the result as the percentage of axonal damage within the inflammatory lesion.

### 2.6. MS patient cohort for M-MDSC blood analysis

31 RR-MS patients fulfilling the 2010-revised McDonald diagnostic criteria [36] who initiated fingolimod treatment were recruited at the Vall d’Hebron Hospital (Barcelona), the Ramón y Cajal Hospital (Madrid) or the Donostia Hospital (San Sebastián). The study was approved by the corresponding local ethics committees, and informed written consent was obtained from all the participants in accordance with the Helsinki declaration. None of the MS patients in this study received corticosteroids in the last 30 days and they were treated with fingolimod (0.5 mg/day) based on current guidelines. In patients who switched treatment from a prior DMT, a minimum washout period of 24 h was established for those who had been receiving interferon-beta (IFN-β) or GA, of 2 months for natalizumab, and of 1 month for azathioprine. All the patients completed 12 months of fingolimod treatment (for demographic and clinical characteristics see Suppl. Table 2).

Neurological evaluation was carried out at baseline, 6 and 12 months after initiation of the fingolimod therapy with additional evaluation in case of relapses. Relapses were defined as a worsening of neurological impairment or the appearance of new symptoms attributable to MS in the absence of fever or infection, lasting at least 24 h and preceded by at least one month of stability [37]. A standardized brain MRI protocol (axial T2, axial T2 FLAIR and T1) was implemented to reliably detect T2 lesions at baseline and new or enlarging lesions at 12 months after initiation of the fingolimod therapy. The transverse T1-weighted sequence was systematically repeated after gadolinium (Gd) injection (0.2 mmol/kg) whenever T2 lesions were demonstrated. Both MRI scans were performed using the same 1.5-T equipment with a slice thickness varying from 2 to 5 mm. Disability was assessed by a trained neurologists using the Kurtzke Expanded Disability Status Scale (EDSS) [38] prior to the initiation of fingolimod therapy and then at 6 and 12 months, and whole blood was collected from these patients at the same time points during follow-up.

### 2.7. Flow cytometry analysis of peripheral blood cells from fingolimod treated patients

Fresh anti-coagulated EDTA blood was obtained by venopunction the same day of the treatment, before the first dose of fingolimod and after 12 months of treatment. Human peripheral blood mononuclear cells (PBMCs) were isolated by Ficoll density gradient centrifugation (GE-171440-02, Merck), collected from the interphase, washed with isolation buffer (2.23 g/L D-glucose, 2.2 g/L sodium citrate, 0.8 g/L citric acid, 0.5% Bovine Serum Albumin in PBS) and further centrifuged at 350g for 10 min at RT. The cell pellet was resuspended in FBS, counted and aliquoted 1:1 in FBS with 20% dimethylsulfoxide (DMSO: Sigma-Aldrich). The samples were then stored long-term in liquid nitrogen at −170°C until use.

Freshly thawed PBMCs (1×10^6^) were washed twice with PBS and stained with Zombie NIR Dye (Biolegend) following the manufacturer’s instructions. After 15 min at RT in the dark, the cells were washed with PBS and recovered at 390g for 5 min. PBMCs were then resuspended in 25 μL of staining buffer and incubated for 10 min at 4°C with Beriglobin (50 μg/mL: CSL Behring) to block the Fc receptors. Subsequently, 25 µL of the antibody mix diluted in staining buffer was added to the cells for 30 min at 4°C in the dark. The antibody panel (Suppl. Table 1) to stain M-MDSCs included anti-CD15-FITC (HI98 clone), anti-CD14-PerCP-Cy^TM^5.5 (Mφ29 clone), anti-CD11b-PE-Cy7 (ICRF44 clone), anti-CD33-APC (WM53 clone) and anti-HLA-DR-BV421 (G46-6 clone: BD Biosciences). After staining, the PBMCs were rinsed with staining buffer at 390g for 5 min at RT, resuspended in 0.1% PFA diluted in PBS and stored at 4°C in the dark until the next day. The samples were then analysed in a FACS Canto II cytometer (BD Biosciences) with the corresponding controls and 100.000 events were recorded in the mononuclear cell (MNC) gate. Compensation controls were obtained by incubating each single antibody with the anti-mouse Ig,k/Negative Control Compensation Particle Set (CompBeads, BD Biosciences). Fluorescence Minus One controls (FMO) were set-up in pooled samples to avoid any variation between the experimental conditions and the data were analysed with FlowJo v.10.6.2 software (FlowJo, LLC-BD Biosciences).

### 2.8. Statistical analysis

The data were analyzed with SigmaPlot version 11.0 (Systat Software Inc.). Age, and gender balance were compared using a Fisher exact test. A student’s *t*-test was used to compare pairs of the different groups of mice/patients with parametric distributions or a Mann-Whitney *U* test was used for non-parametric distributions. A Spearman test was carried out for the correlation analyses. A ROC curve analysis was performed to determine the discriminative ability of the different clinical parameters scored and to determine the threshold of M-MDSCs, with the highest sensitivity and specificity to separate responder from non-responder mice. In addition, treatment efficacy was also determined by a ROC curve analysis. Finally, a Cox regression analysis was performed to determine the probability of reaching the population median of the clinical parameters presented by the vehicles as a function of different categorical and non-categorical variables. The minimum statistical significance was set at p<0.05: *p<0.05, **p<0.01, ***p<0.001.

## 3. Results

### 3.1. Individualized fingolimod treatment accelerates disease recovery in EAE

Fingolimod has been seen previously to improve the clinical course of EAE [39, 40], although in all these earlier studies fingolimod treatment was carried out on randomly assigned groups of mice with diverse clinical scores, without taking into account the disease severity of each individual animal. In order to better approximate to the clinical situation of MS patients, we used an individualized treatment protocol that started at the specific point of disease onset for each animal. Differences between the experimental groups were observed from 1 dpo, which became constantly significant from 4 dpo onwards. In general terms, the individualized fingolimod treatment significantly reduced the maximum clinical score of each animal, enhancing their recovery at a higher and faster rate (the percentage recovery and recovery index, respectively). In addition, a decrease in their residual score and reduced total accumulated score was observed relative to the EAE-Veh animals (Figure 1A, Table 1). The amelioration in the clinical signs observed in mice administered fingolimod was paralleled by a reduction in demyelination: Demyelinated area-EAE-Vehicle: 59,796.97 ± 22,444.54, EAE-FTY720: 22,168.25 ± 15,461.25, p<0.043); % of the demyelinated area within the white matter (EAE-Veh: 6.26 ± 2.30, EAE-FTY720; 2.14 ± 1.44, p<0.042: Figure 1B,C and F); and % of axonal damage within the demyelinated area (EAE-Veh: 0.98 ± 0.22, EAE-FTY720: 0.43 ± 0.12, p<0.045: Figure 1D, E and G). Moreover, the individualized fingolimod treatment also dampened the abundance of splenic CD3^+^, CD8^+^ and CD3^+^CD25^+^ T lymphocytes (Table 1).

**Figure 1:**
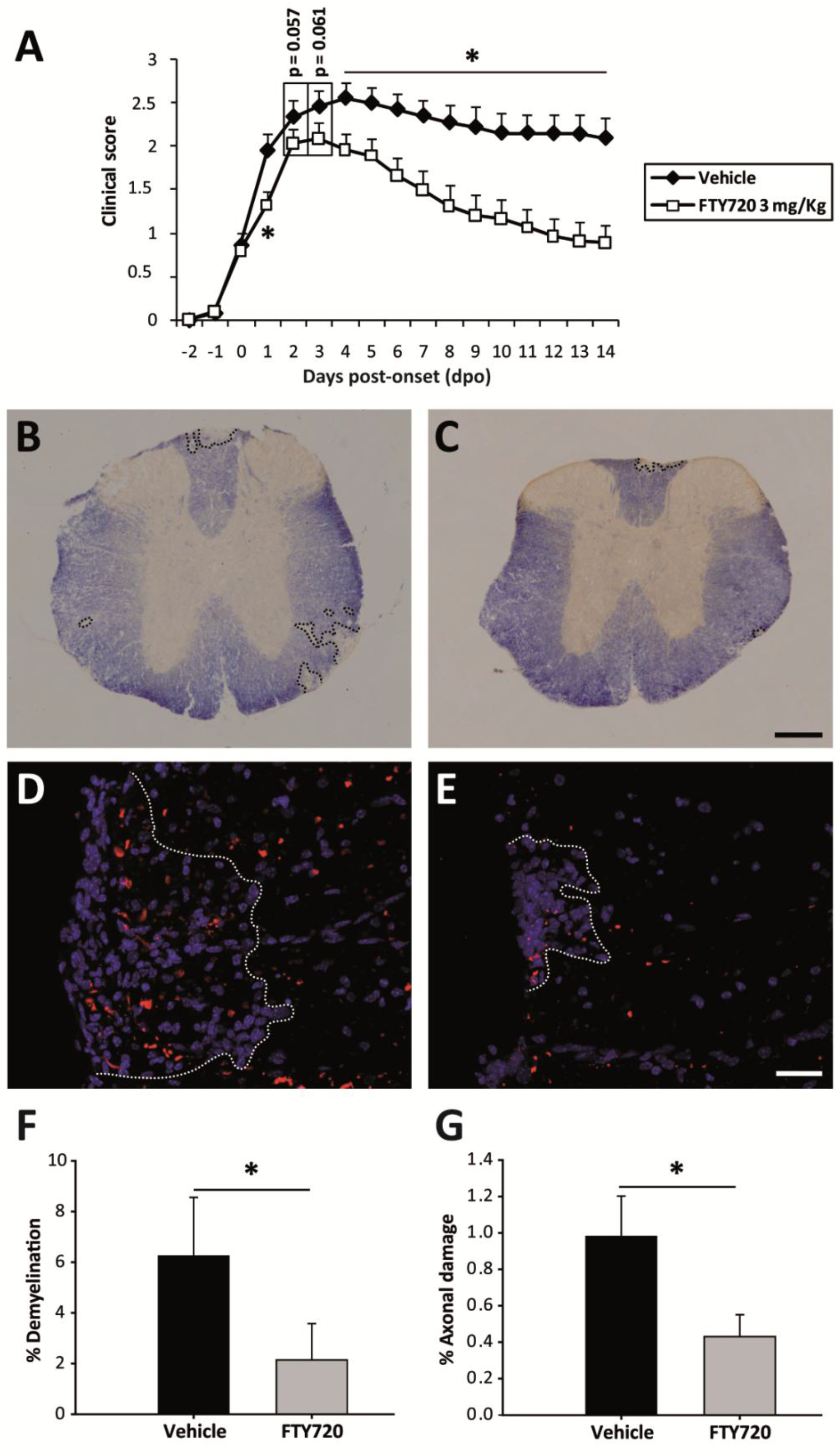
Individualized fingolimod treatment accelerates disease recovery and decrease the histopathological damage in EAE mice. **A:** Individualized fingolimod treatment from disease onset significantly reduced the clinical course and severity of EAE in mice (EAE-Veh N = 19; EAE-FTY720 N =18). **B, C**: Panoramic images of the spinal cord of one EAE-Veh (**B**) and one EAE-FTY720 (**C**) mouse stained with EC (myelin, blue). Dashed lines define the demyelinated area. **D, E**: Detailed view of SMI-32 staining (axonal damage, red) in infiltrated areas (delimitated by dashed lines) of the spinal cord of EAE-Veh (**D**) and EAE-FTY720 (**E**) mice (nuclei are stained with Hoechst). **F-G**: Quantification of the relative demyelination within the white matter or the axonal damage within the infiltrated area in the histological sub-cohort of EAE mice (EAE-Veh N = 9, EAE-FTY720 N =9). Scale bar: **B-C**= 250 µm; **D-E** = 50 µm.

**Table 1.** Comparison of the clinical parameters and splenic T cell populations in EAE mice following individualized treatment with fingolimod or vehicle.

|  |  | **Vehicle** | **FTY720** | **P value** |
| --- | --- | --- | --- | --- |
| **CLINICAL PARAMETERS** | **Maximum score** | 3<br>(2.25 – 3) | 2.38<br>(1.18 – 2.75) | **<0.05** |
|  | **dpi peak** | 15<br>(14 – 17) | 15<br>(14 – 15.25) | *ns* |
|  | **Severity Index** | 0.75<br>(0.50 – 1.00) | 0.625<br>(0.33 – 0.75) | *ns* |
|  | **Peak accumulated score** | 6.50<br>(5.63 – 9.00) | 5.69<br>(3.68 – 6.81) | *ns* |
|  | **Total accumulated score** | 37.25<br>(23.00 – 42.38) | 18.88<br>(12.59 – 27.16) | **< 0.01** |
|  | **% Recovery** | 25.00<br>(12.50 – 33.30) | 72.73<br>(53.41 – 80.24) | **< 0.01** |
|  | **Recovery Index** | 0.093<br>(0.04 – 0.17) | 0.16<br>(0.10 – 0.21) | **< 0.05** |
|  | **Residual score** | 2.25<br>(1.00 – 2.63) | 0.50<br>(0.50 – 0.88) | **< 0.01** |
| **SPLEEN** | **% CD3<sup>+</sup>/Total blood cells** | 15.15 ± 0.97 | 11.06 ± 0.93 | **< 0.01** |
|  | **% CD4<sup>+</sup>/Total blood cells** | 9.11 ± 0.64 | 7.75 ± 0.63 | *ns* |
|  | **% CD8<sup>+</sup>/Total blood cells** | 4.98 ± 0.36 | 2.79 ± 0.28 | **<0.001** |
|  | **% CD3<sup>+</sup>CD25<sup>+</sup>/Total blood cells** | 1.32 ± 0.08 | 1.09 ± 0.07 | **< 0.05** |
|  | **% CD3<sup>+</sup>CD69<sup>+</sup>/Total blood cells** | 1.77 ± 0.12 | 1.86 ± 0.10 | *ns* |

The data are expressed as the median (IQR) for the clinical parameters and the mean ± SEM for the splenic parameters. The results of applying a Student t-test or a non-parametric Mann-Whitney test are shown: dpi, day post-immunization; IQR, interquartile range; ns, non-significant.

In summary, the individualized follow-up in each mouse from the point of disease onset appeared to be a good strategy to explore the differential efficacy of fingolimod treatment in EAE.

### 3.2. The abundance of Ly-6C^hi^cells in peripheral blood is related to a milder long term EAE course in fingolimod administered mice

To establish the M-MDSC values that might predict a future response to fingolimod in MS, we explored the relationship between their abundance in the blood and the future short and long term clinical course in fingolimod administered EAE mice. Highly immunosuppressive M-MDSCs in the spleen and spinal cord of EAE mice, as well as circulating M-MDSCs, are phenotypically indistinguishable from CD11b^+^Ly-6C^hi^Ly-6G^-/low^ inflammatory monocytes, also called Ly-6C^hi^ cells [18,20,21,41,42]. Interestingly, high levels of splenic Ly-6C^hi^ cells at the peak of EAE have been directly associated with a milder disease course [24, 43]. Hence, to explore whether M-MDSCs could be used a predictive tool for fingolimod-induced reduction in the severity of the clinical course of EAE, we first investigated the correlation between the abundance of Ly-6C^hi^ cells in the peripheral blood at disease onset and different short- and long-term clinical parameters of the clinical course of EAE. As in the case of EAE-Veh, the higher proportion of Ly-6C^hi^ cells relative to the total blood content was directly related to a milder EAE disease course: lower maximum score at peak, lower total accumulated and residual clinical scores, and stronger relative disease recovery in EAE-FTY720 mice (Figure 2A-D). These data pointed to the quantification of Ly-6C^hi^ cells as an interesting parameter in the search for biomarkers that link disease severity reduction after fingolimod treatment.

**Figure 2.**
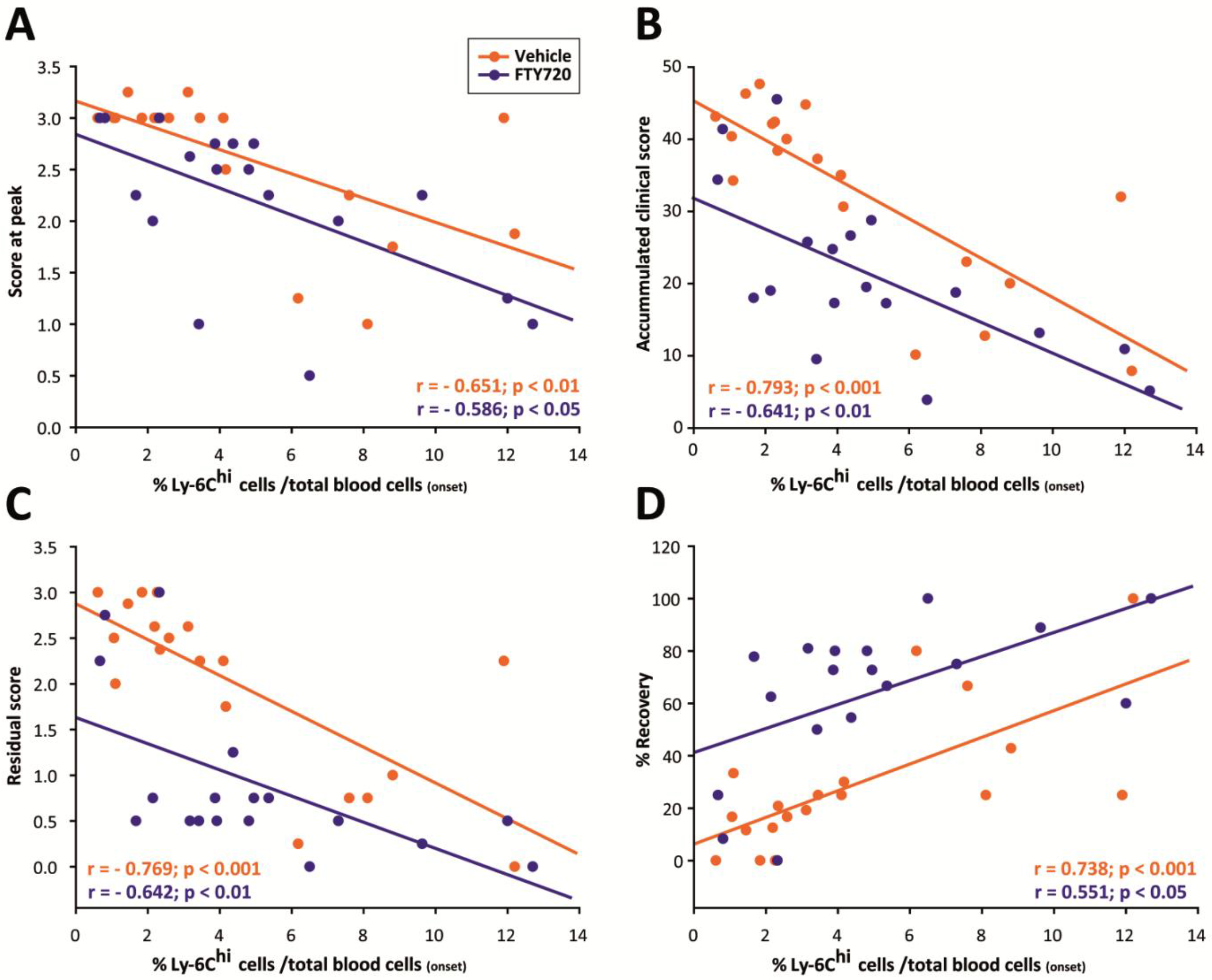
Ly-6Chi cell abundance at EAE onset is related to milder short- and long-term disease courses. **A-D:** The amount of circulating Ly-6C^hi^ cells at disease onset is inversely correlated with the peak score (**A**), the total accumulated clinical score (**B**) and the residual score (**C**), and it is directly correlated with the % recovery (**D**) in both EAE-Veh and EAE-FTY720 mice.

To check whether the initial randomized distribution of EAE mice influenced the long-term effects of fingolimod, we compared the clinical and immunological baseline conditions between the two experimental groups. The day of onset and the clinical score at that point did not differ between these groups: day at onset (EAE-Veh 13 dpi-Inter Quartile Range–IQR 12–14), EAE-FTY720: 13 dpi IQR 12–13, p = 0.789); and clinical score at onset (EAE-Veh: 0.75 IQR 0.50-0.75), EAE-FTY720: 0.75 IQR 0.5-1.0, p = 0.763). Importantly, no major differences were reported in the abundance of Ly-6C^hi^ cells between the animals treated with vehicle or fingolimod, neither relative to the total number of blood cells or in the myeloid subset (Figure 3A-B). Moreover, the Ly-6C^hi^ cells from both animal groups showed a similar mean fluorescence intensity (MFI) in terms of their activation or the differentiation markers expressed prior to vehicle or fingolimod administration (Figure 3C-E). Therefore, these data clearly indicated that the content and characteristics of the circulating Ly-6C^hi^ cells was not biased between the two groups of animals prior to treatment.

**Figure 3.**
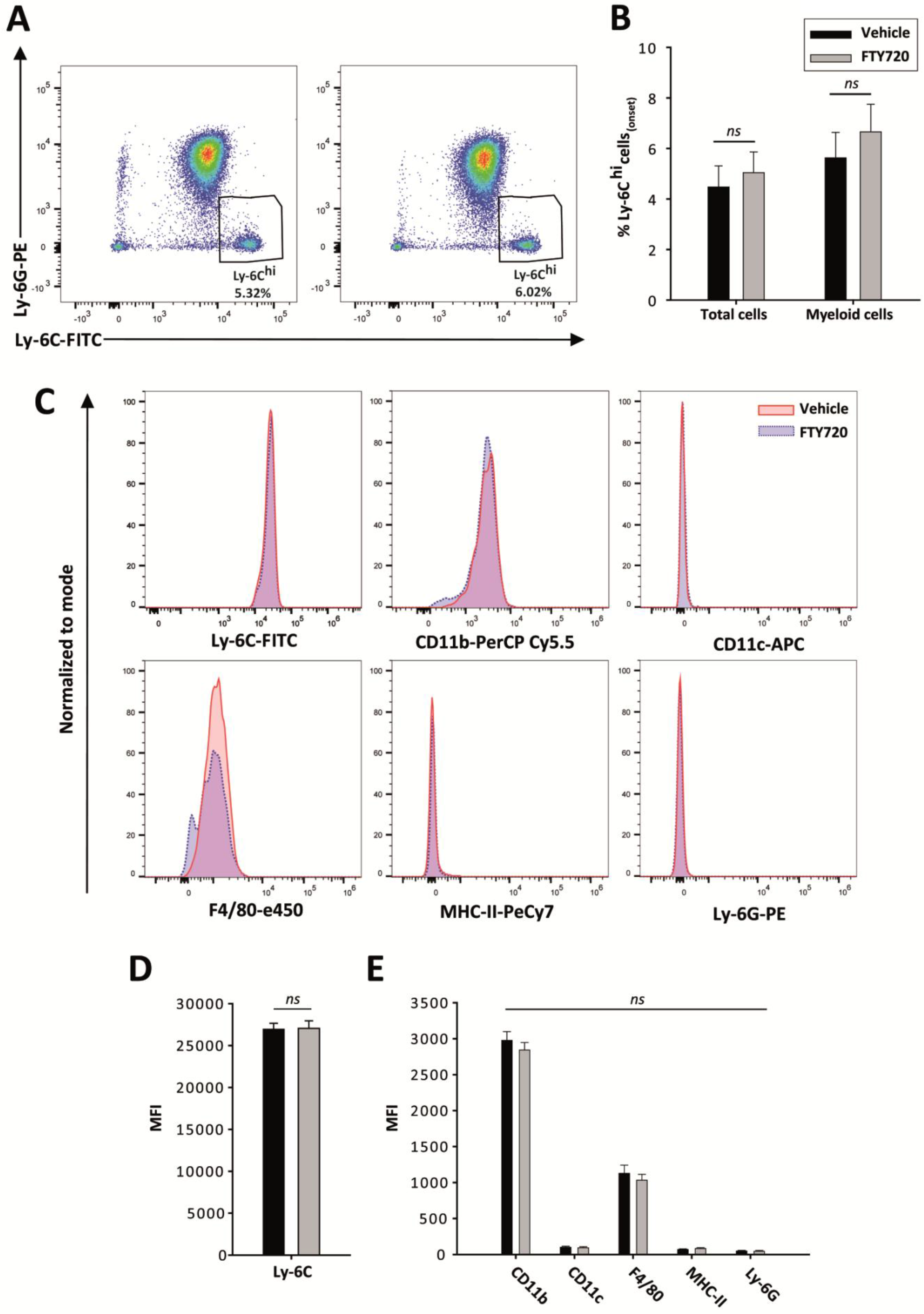
Ly-6C^hi^ cells are homogeneously represented in EAE-Vehicle and EAE-FTY720 mice before treatment. **A:** Representative plots of Ly-6C^hi^ cells from EAE mice that will be treated with vehicle (left panel) or fingolimod (right panel). **B:** The abundance of Ly-6C^hi^ cells relative to the total cells or to the myeloid component at disease onset was similar between both experimental study groups prior to the beginning of treatment. **C:** Representative histograms from the main phenotypic markers of circulating Ly-6C^hi^ cells at disease onset. **D-E:** The MFI of the different markers of Ly-6C^hi^ cells did not differ prior to onset of treatment.

We also looked for differences in the abundance of circulating T cells, the main targets for both fingolimod and MDSCs, prior to vehicle/fingolimod administration. No differences were detected in CD3^+^ or CD8^+^ T lymphocytes, although there was a slightly higher percentage of CD4^+^ T lymphocytes in animals that then received fingolimod (Table 2). To check whether the difference in CD4^+^ T lymphocytes might reflect an imbalance in the immunological environment, we compared the Ly-6C^hi^/T lymphocyte ratio between the two experimental groups. No differences were detected in the relative abundance of Ly-6C^hi^ cells and CD3^+^, CD4^+^ or CD8^+^ T lymphocytes between mice administered fingolimod or the vehicle alone, downplaying the significance of the differences at baseline after considering the inflammatory microenvironment as a whole. These results clearly showed that the therapeutic effect of fingolimod observed is not related to a clinical or immunological bias associated with the individualized treatment of EAE.

**Table 2.** Immunological cells in the blood of EAE mice at disease onset

|  | *Vehicle* | *FTY720* | *P value* |
| --- | --- | --- | --- |
| % $CD3^+$ /Total blood cells | 12.94 ± 1.32 | 16.77 ± 1.63 | *ns* |
| % $CD4^+$ /Total blood cells | 8.09 ± 0.77 | 10.91 ± 1.03 | **<0.05** |
| % $CD8$ /Total blood cells | 3.32 ± 0.30 | 4.42 ± 0.51 | *ns* |
| % $Ly-6C^{hi}$ /Total blood cells: % $CD3^+$ | 0.47 ± 0.13 | 0.39 ± 0.10 | *ns* |
| % $Ly-6C^{hi}$ /Total blood cells: % $CD4^+$ | 0.72 ± 0.19 | 0.58 ± 0.15 | *ns* |
| % $Ly-6C^{hi}$ /Total blood cells: % $CD8^+$ | 1.83 ± 0.55 | 1.68 ± 0.50 | *ns* |
| % $Ly-6C^{hi}$ /Myeloid cells: % $CD3^+$ | 0.58 ± 0.16 | 0.50 ± 0.13 | *ns* |
| % $Ly-6C^{hi}$ /Myeloid cells: % $CD4^+$ | 0.89 ± 0.23 | 0.75 ± 0.18 | *ns* |
| % $Ly-6C^{hi}$ /Myeloid cells: % $CD8^+$ | 2.25 ± 0.65 | 2.15 ± 0.62 | *ns* |

The data are expressed as the mean ± SEM. The result of the Student t-test or a non-parametric Mann-Whitney test is shown: *ns*, not significant.

### 3.3. Circulating Ly-6C^hi^ cells represent an excellent biomarker to predict the responsiveness to fingolimod in EAE mice

Although fingolimod was therapeutically effective in most of the EAE-FTY720 mice (responder mice), 3 of the 18 mice showed a clinical course indistinguishable from the control EAE-Veh mice, i.e. non-responders (Figure 4A-C). To quantify this effect, we considered non-responder mice as those with a similar or worse clinical level than the median of the EAE-Veh group. Thus, EAE-FTY720 non-responders were defined as those with a maximum clinical score ≥3, with a clinical recovery score ≤25 %, a recovery of the clinical score between the peak and the end of the follow-up ≤0.5 and a residual clinical score ≥2.25. Following this distribution, EAE-FTY720 responder mice exhibited a significantly milder disease course than EAE-Veh or EAE-FTY720 non-responders, which in turn showed no difference between them (Table 3). Indeed, similarities between EAE-Vehicle mice and EAE-FTY720 non-responders were also observed in the splenic immunological status at the end of the follow-up. Interestingly, only responder mice had a significantly smaller proportion of splenic CD3^+^, CD4^+^, CD8^+^ and CD3^+^CD25^+^T lymphocytes (Table 3).

**Figure 4.**
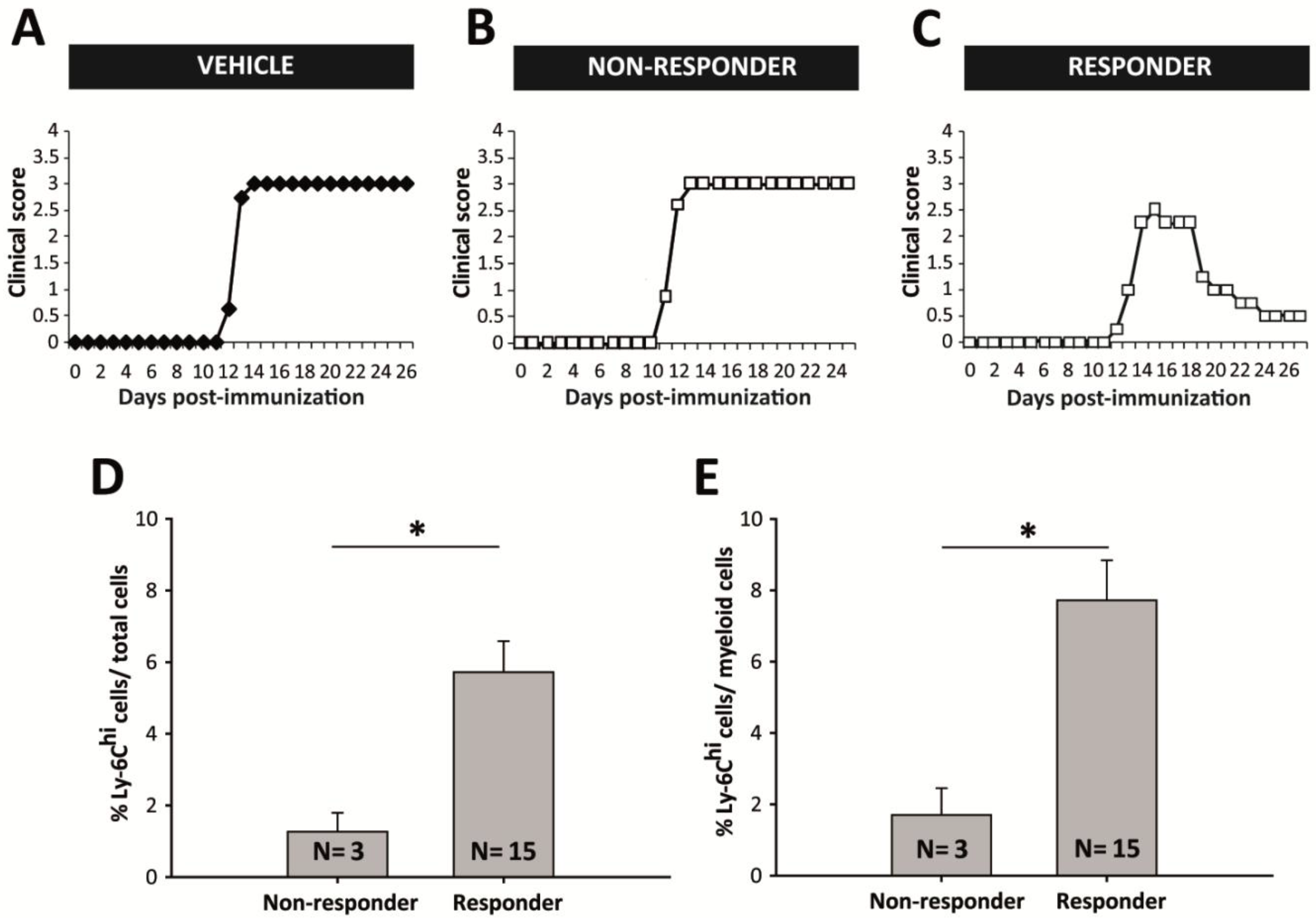
Circulating levels of Ly-6C^hi^ cells are higher in fingolimod responder EAE mice. **A-C:** Graphic representation of the clinical course of one representative EAE-Veh (**A**: Ly-6C^hi^cells/total blood cells = 2.26% or Ly-6C^hi^ cells/myeloid cells= 3.15%), one non-responder (**B**: Ly-6C^hi^cells/total blood cells = 2.32%; Ly-6C^hi^cells/myeloid cells = 3.20%) and one responder EAE-FTY720 mouse (**C**: Ly-6C^hi^cells/total blood cells = 4.81% or Ly-6C^hi^cells/myeloid cells= 6.02%). EAE-Veh animals developed a severe clinical course undistinguishable from non-responder EAE-FTY720 mice. **D-E**: Responder EAE-FTY720 mice (N = 3) had a much higher abundance of circulating Ly-6C^hi^cells among the total blood cells (**D**) or the myeloid component (**E**) than non-responder EAE-FTY720 mice (N = 15) prior to initiating treatment.

**Table 3.** Comparison of the clinical parameters and splenic T cell populations in EAE-Vehicle, and EAE-FTY720 responder and non-responder mice.

|  |  | EAE-Vehicle | EAE-FTY720 |  | P value |
| --- | --- | --- | --- | --- | --- |
|  |  |  | Non-responder | Responder |  |
| CLINICAL PARAMETERS | Maximum score | 3<br>(2.25-3) | 3<br>(3-3) | 2.25*<br>(1.25-2.63) | <0.01 |
|  | dpi peak | 15<br>(14-17) | 15<br>(13-15) | 15<br>(14-16) | ns |
|  | Severity Index | 0.75<br>(0.50-1) | 0.75<br>(0.60-1) | 0.63<br>(0.33-0.69) | ns |
|  | Peak accumulated score | 6.5<br>(5.63-9) | 7<br>(6.50-9.75) | 4.5*<br>(3-6.63) | <0.05 |
|  | Total accumulated score | 37.25<br>(23-45.38) | 41.38<br>(34.38-45.50) | 18*<br>(10.90-24.75) | <0.001 |
|  | % Recovery | 25<br>(12.5-33.30) | 8.33<br>(0-25) | 75*<br>(62.50-80.95) | <0.001 |
|  | Recovery Index | 0.09<br>(0.04-0.17) | 0.04<br>(0-0.09) | 0.17*<br>(0.13-0.22) | <0.01 |
|  | Residual score | 2.25<br>(1-2.63) | 2.75<br>(2.25-3) | 0.5*<br>(0.5-0.75) | <0.001 |
| SPLEEN | % CD3<sup>+</sup>/Total blood cells | 15.15 ± 0.97 | 15.68 ± 3.27 | 10.13± 0.77* | <0.01 |
|  | % CD4<sup>+</sup>/Total blood cells | 9.11 ± 0.64 | 10.51 ± 2.17 | 7.20 ± 0.56* | <0.05 |
|  | % CD8<sup>+</sup>/Total blood cells | 4.98 ± 0.36 | 4.47 ± 0.91 | 2.46 ± 0.21* | <0.001 |
|  | % CD3 CD25<sup>+</sup>/Total blood cells | 1.32 ± 0.08 | 1.33 ± 0.25 | 1.04 ± 0.07* | <0.05 |
|  | % CD3 CD69<sup>+</sup>/Total blood cells | 1.77 ± 0.12 | 2.24 ± 0.31 | 1.78 ± 0.10 | ns |

The data are expressed as the median (IQR). One Way ANOVA or a non-parametric ANOVA on Ranks tests are shown: dpi, day post-immunization; ns, not significant; IQR, interquartile range. \*represents significantly different groups.

When the level of circulating Ly-6C^hi^ cells in the blood prior to fingolimod administration was analyzed, EAE-FTY720 non-responder mice had a remarkable lower abundance of these cells within the whole blood cells than EAE-FTY720 responders prior to treatment initiation (Figure 4D, E). Importantly, Ly-6C^hi^ cell levels had excellent discriminatory power to assess the risk of being responder or non-responder based on the median value of the maximum peak clinical score, the percentage recovery of the clinical score and the residual clinical score (AUC 0.956, 95% CI –confidence interval-0.853-1.058; p<0.05). In all cases, we established 2.75% of Ly-6C^hi^ cells within the whole leukocyte population at disease onset as the cut-off value indicative of a future therapeutic responsiveness in EAE mice (100% specificity, 95% C.I. 29.24%-100%; 86.67% sensitivity, 95% C.I. 59.54%-98.34%; Figure 5A-B). By contrast, the mere abundance of this cell type was not able to discriminate between responder and non-responder EAE-FTY720 mice based on the median accumulated clinical score at the end of the experiment (AUC 0.875, 95% CI 0.703-1.047; p =0.092) or on the recovery index (AUC 0.667, 95% CI 0.305-1.028; p = 0.374; Figure 5C-F).

**Figure 5.**
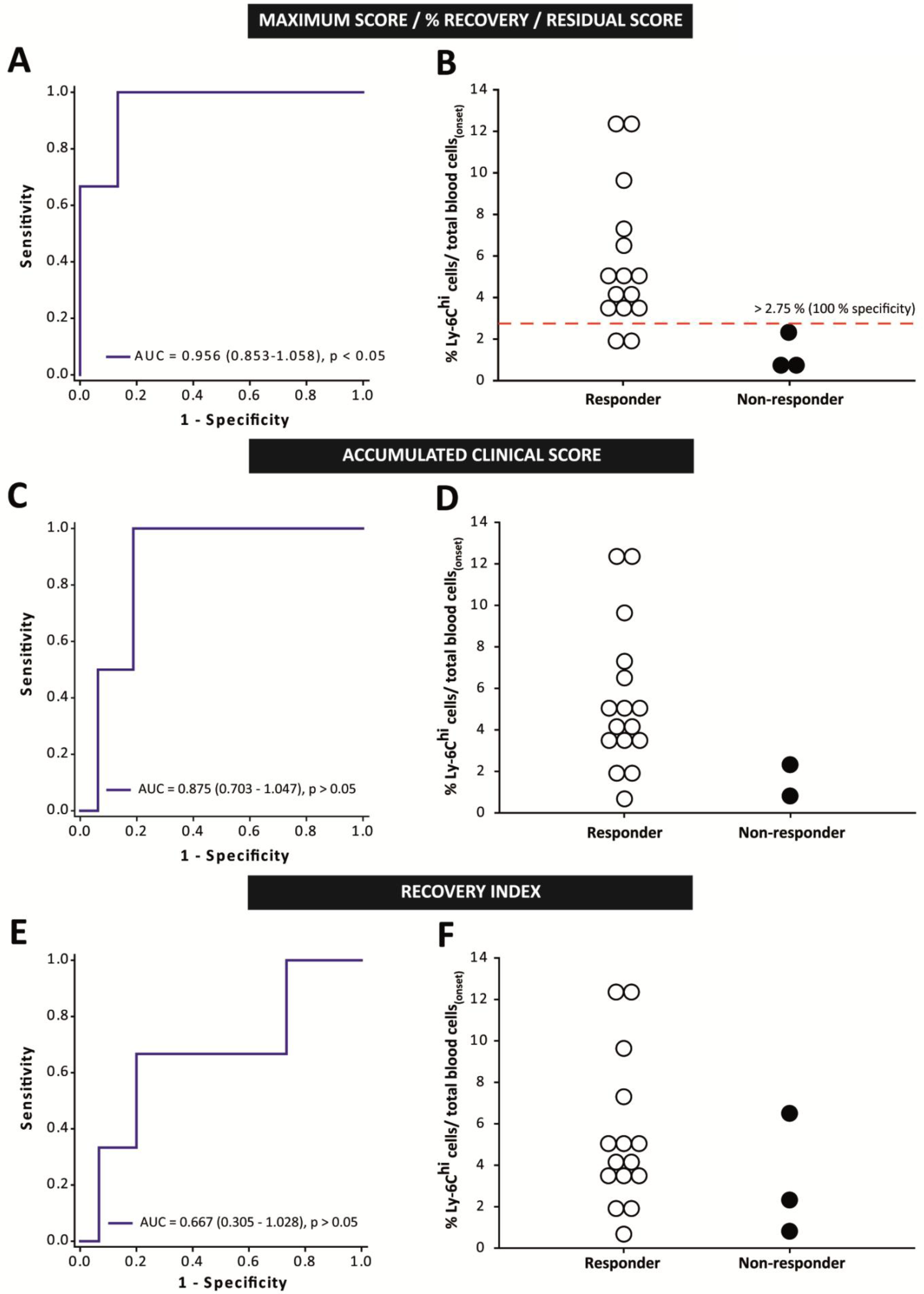
Ly-6C^hi^ cell content is an excellent biomarker to predict responsiveness to fingolimod in EAE mice. **A:** ROC curve analysis showing that Ly-6C^hi^ cell content presented the highest discriminatory ability for responder and non-responder EAE mice classified using the median of maximum clinical score, the % recovery or the residual score. **B:** Cut-off value for the Ly-6C^hi^cells/total blood cells to predict good responsiveness to fingolimod. **C-F:** The level of Ly-6C^hi^ cells could not discriminate between responder and non-responder EAE-FTY720 mice based on the median of the total accumulated clinical score (**C-D**) or the recovery index (**E-F**). AUC,area under the curve (N= 18 mice).

In summary, our data clearly indicated that circulating Ly-6C^hi^ cell content at the onset of EAE is an extraordinary tool to determine future therapeutic responsiveness of EAE mice to fingolimod treatment in terms of the maximum affectation, the extent of recovery and the residual score achieved by each individual animal.

### 3.4. Circulating Ly-6C^hi^cells constitute a good biomarker of fingolimod treatment efficacy in EAE

As in clinical practice, examination of EAE responder mice yielded high variability in the efficacy demonstrated by fingolimod treatment in each individual mouse. In fact, our data revealed that the higher the number of circulating Ly-6C^hi^cells prior to treatment, the greater the difference in the maximum score achieved by EAE-FTY720 mice relative to the median score of the EAE-Veh mice (Figure 6A). Hence, Ly-6C^hi^ cells may be a putative biomarker not only of responsiveness to fingolimod treatment but also, to the individual degree of drug efficacy.

**Figure 6.**
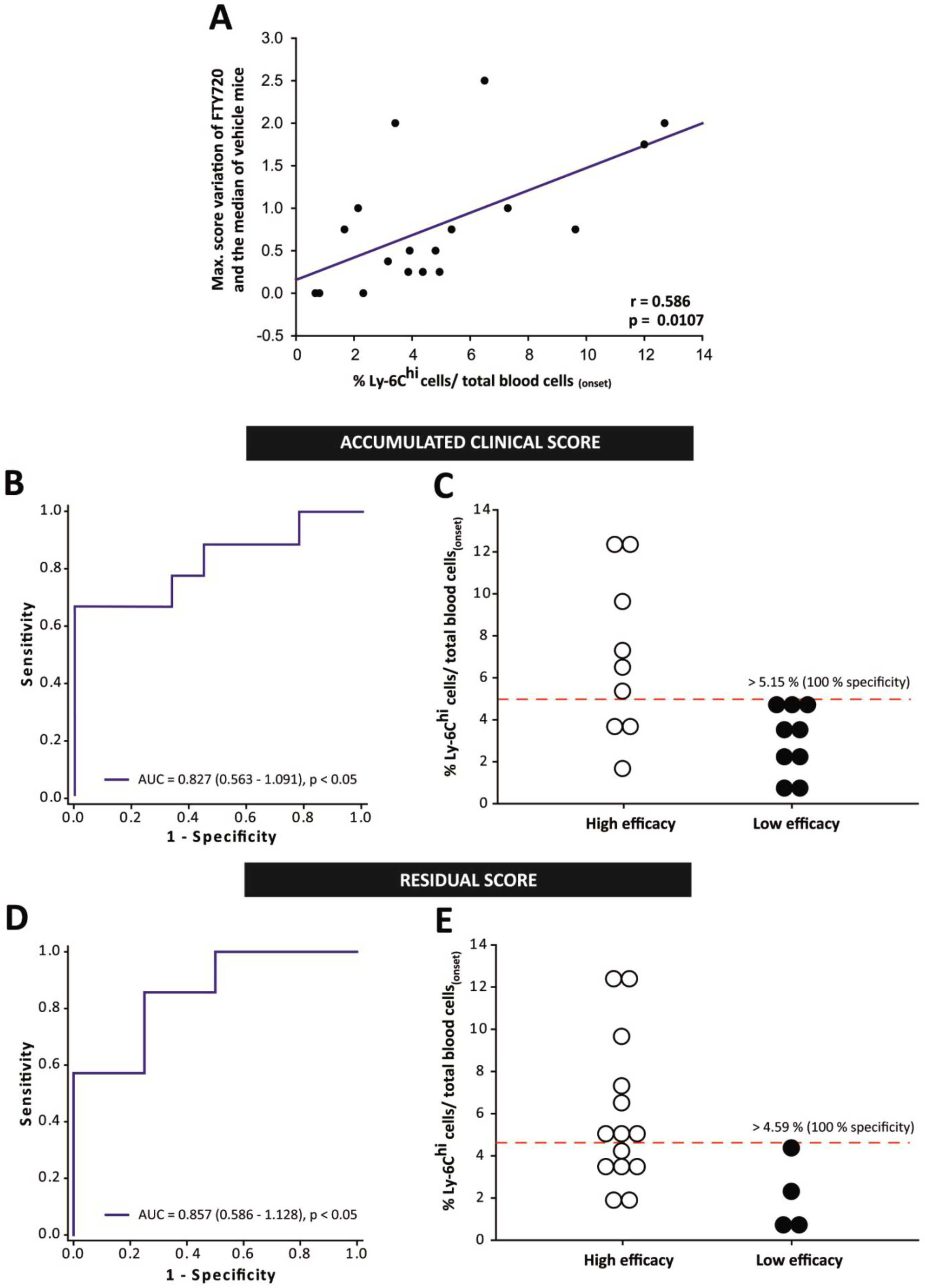
Ly-6C^hi^ cells are a good biomarker for the efficacy of fingolimod treatment in EAE. **A:** The higher the Ly-6C^hi^cell content at onset, the greater the difference in the maximum score reached by EAE-FTY720 mice compared to the median of the EAE-Veh group. **B,D:** ROC curve analysis showing that Ly-6C^hi^ cells had a good capacity to discriminate high efficacy as opposed to low efficacy EAE mice through the median accumulated clinical score (**B**) and the residual score (**D**). **C,E:** Cut-off values for the Ly-6C^hi^cells/total blood cells to predict those EAE mice that will present a higher total accumulated or a lower residual score than the median of EAE-FTY720 mice at the end of the experimental evaluation.

To assess this phenomenon, we classified the effect of fingolimod in each EAE mouse as high or low based on the median of the EAE-FTY720 group as a whole. Fingolimod treatment was considered to be high efficacy when an EAE-FTY720 mouse had a maximum clinical score <2.5, a total accumulated clinical score <19, clinical recovery >72.73%, a recovery index >0.16 or a residual clinical score <1.0. The abundance of Ly-6C^hi^ cells prior to fingolimod treatment could not discriminate between animals prone to present high/low treatment efficacy based on the median of the maximum clinical score (AUC 0.765, 95% CI 0.524-1.007; p = 0.058), the % recovery of the clinical score (AUC 0.700, 95% 0.450-0.950; p= 0.155), or the recovery index (AUC 0.741, 95% CI 0.491-0.990; p = 0.085). Interestingly, Ly-6C^hi^ cell level was seen to be an important parameter to distinguish EAE mice in which fingolimod acts with high efficacy based on the median value of the total accumulated clinical score (Figure 6B) and of the residual score at the end of fingolimod administration (Figure 6D). In fact, having more than 5.15% (100% specificity, 99% C.I. 55.50%-100%; 66.67% sensitivity, 99% C.I. 21.91%-95.84%; Figure 6C) or 4.59 % (100% specificity, 95% C.I. 26.59%-100%; 57.14% sensitivity, 95% C.I. 22,34-87,33%; Figure 6E) of circulating Ly-6C^hi^ cells at disease onset were valid cut-off points in EAE mice to distinguish those prone to experience high efficacy of fingolimod treatment in terms of the total accumulated clinical score or residual score, respectively.

In summary, the proportion of peripheral blood Ly-6C^hi^ cells at disease onset appeared to be a good biomarker of fingolimod efficacy, whereby the higher the level of these cells the less severe the disease after fingolimod treatment.

### 3.5. The responsiveness and efficacy to fingolimod treatment in EAE is exclusively influenced by the level of circulating Ly-6C^hi^ cells

The existence of differences in the responsiveness and efficacy to fingolimod could be the result of the sum of different clinical and immunological factors other than Ly-6C^hi^ cell levels. To test this hypothesis, we performed a Cox proportional hazard regression analysis to determine the likelihood of EAE mice achieving a maximum clinical score ≥3 (median of the EAE-Veh mice, i.e. paraplegia) or a residual score <2 as a function of the following variables found at disease onset: dpi; clinical score; the abundance of Ly-6C^hi^ cells; CD3^+^, CD4^+^, CD8^+^ T cells; the ratios between Ly-6C^hi^ cells/lymphocyte subsets, and some other myeloid cell subsets, such as neutrophils (CD11b^+^Ly-6C^int^Ly-6G^high^) or dendritic cells (CD11b^+^CD11c^+^); and the experimental animal group, i.e. treated or not with fingolimod. Importantly, Ly-6C^hi^ cell abundance at disease onset and the absence of fingolimod treatment were the only two variables associated with a lower risk of reaching paraplegia or completing the treatment with a residual clinical score <2 (p <0.05 in both cases). A high abundance of Ly-6C^hi^ cells at disease onset was a clear protective factor (HR = 0.0949, 95% C.I. 0.0106 −0.851) while the absence of fingolimod treatment was a prominent risk factor for reaching paraplegia (HR = 11.195, 95% C.I. 1.212-103.397). On the other hand, a high abundance of Ly-6C^hi^ cells at disease onset increased the likelihood of completing the treatment with little clinical sequelae (HR = 1.894, 95% C.I. 1.238-2.898), whereas not being treated with fingolimod prevented such an outcome (HR = 0.149, 95% C.I. 0.0358-0.623). All these data clearly indicate that the abundance of Ly-6C^hi^ cells is a good and specific biomarker for fingolimod treatment efficacy in the EAE model.

### 3.6. M-MDSC abundance is independent of the fingolimod to treat MS patients’ characteristics at baseline

To establish a parallel with the observations in the EAE mouse model, we carried out a translational study in 31 MS patients (demographic data are shown in Suppl. Table 2) to assess whether circulating M-MDSC abundance at baseline may be related to the future response of MS patients to fingolimod. We first assessed whether the M-MDSC content of the mononuclear cell fraction (M-MDSCs/MNCs) at baseline was independent of the patients’ age (r= 0.0649, p= 0.726), EDSS (r= −0.102, p= 0.584), or number of Gd^+^ enhancing lesions (r = 0.199, p = 0.279). Switching treatment from natalizumab (Ntz) to fingolimod has been associated with a risk of MS reactivation [44–46]. In order to avoid misinterpretation of the M-MDSC abundance due to the presence of patients with a very active immune response, we discriminated patients who switched treatment from Ntz (Ntz-patients, N=11) from those who were naïve or who had previously received another first line treatment (Non-Ntz patients, N= 20). Interestingly, the level of M-MDSCs at baseline was similar between Non-Ntz and Ntz patients (Non-Ntz 1.23 ± 0.30, Ntz 1.22 ± 0.66; p= 0.397). In fact, the M-MDSCs in both Ntz and Non-Ntz patients remained independent of the baseline parameters: age (Non-Ntz r=-0.125, p=0.594; Ntz r=0.397, p=0.210); EDSS (Non-Ntz r=-0.144, p=0.538; Ntz r=0.0961, p= 0.755), number of Gd^+^ enhancing lesions (Non-Ntz r=0.179, p=0.444; Ntz r=0.100, p=0.755). Therefore, our data indicated that M-MDSC abundance at baseline was very clearly independent of demographics at and switching from Ntz treatment.

### 3.7. M-MDSC abundance at baseline is greater in fingolimod responder patients

To establish whether the abundance of M-MDSCs might be related to the future response to fingolimod at 12 months, the baseline M-MDSC level was correlated to the different criteria considered to define non-evidence of disease activity (NEDA-3), i.e. no new relapses; absence of confirmed disability progression (CDP), understood as 1 point increase in the EDSS for patients with a basal EDSS ≤ 5 (1.5 points when baseline EDSS was 0) or of 0.5 points in patients with basal EDSS >5; and no increase in brain MRI activity [a composite measure of any new T2 and/or gadolinium-enhancing lesions (Gd+ lesions)] relative to the baseline MRI [14].

After 12 months of treatment, 74.19% of patients had not experienced relapses, with a significantly lower annual relapse rate (ARR) than the year before starting fingolimod treatment (1.10 ± 0.18 *vs* 0.32 ± 0.11; p<0.001). Despite the fact that 77.42% of patients maintained or decreased their EDSS after treatment, no significant differences were observed in the EDSS score before and after fingolimod treatment [3.0 (2.0-4.0) *vs* 2.5 (2.0-4.0); p=0.603]. Finally, the proportion of patients with no new T2 lesions after 12 months of fingolimod treatment was 61.29%, whereas 87.10% of patients did not present new Gd^+^ lesions after 12 months of follow-up. However, M-MDSC abundance at baseline was not related to the future clinical course at 12 months after treatment (Suppl. Table 3).

To address the ability of M-MDSCs to discriminate between MS patients who will respond distinctly to fingolimod, “responder” MS patients were considered as those who met all the NEDA-3 criteria (R-MS_NEDA-3_), whereas “non-responder” MS patients were those who met two or less NEDA-3 criteria at the end of the follow-up period (NR-MS_NEDA-3_). Accordingly, 13 of the 31 patients (41.94%) were considered as R-MS_NEDA-3_. No relationship was found between the abundance of M-MDSCs and the different clinical parameters found after 12 months follow-up in R-MS_NEDA-3_ (Suppl. Table 3) and hence, the percentage of M-MDSCs at baseline was not significantly different between the R-MS_NEDA-3_ and NR-MS_NEDA-3_ subgroups (R-MS_NEDA-3_ 1.54 ± 0.56, NR-MS_NEDA-3_ 1.01 ± 0.32; p=0.447).

In a second approximation, we restricted the concept of “responder” MS patients to those the so-called clinical response (R-MS_CR_), i.e. MS patients with no new relapses and no CDP after 12 months of treatment [47, 48]. In this sense, 20 of the 31 patients (64.52%) were considered as R-MS_CR_. Again, no relationship was found between the abundance of M-MDSCs and the different clinical parameters found after 12 months follow-up in R-MS_CR_ (Suppl. Table 3). Moreover, the percentage of M-MDSCs at baseline was not significantly different between the R-MS_CR_ and NR-MS_CR_ subgroups (R-MS_CR_ 1.44 ± 0.39, NR-MS_CR_ 0.85 ± 0.45; p=0.094).

In a next step, we restricted the concept of “responder” MS patients (R-MS) to those showing any good therapeutic response (GTR), i.e. MS patients who meet 2 or more of the NEDA-3 criteria after 12 months of treatment [14, 15]. Accordingly, “non-responder” MS patients (NR-MS) were those who met one or none NEDA-3 criteria. In this sense, 25 out of 31 patients (80.61%) were classified as R-MS. Again, no relationship was established between the abundance of M-MDSCs and the clinical parameters after a 12 month follow-up within the R-MS group (Suppl. Table 3). Importantly, R-MS patients did have a significantly higher percentage of M-MDSCs than NR-MS (R-MS 1.48 ± 0.35, NR-MS 0.16 ± 0.06; p<0.05: Figure 7A-B). As such, the abundance of M-MDSCs did appear to be a useful tool to identify MS patients with a suboptimal therapeutic response to fingolimod.

**Figure 7:**
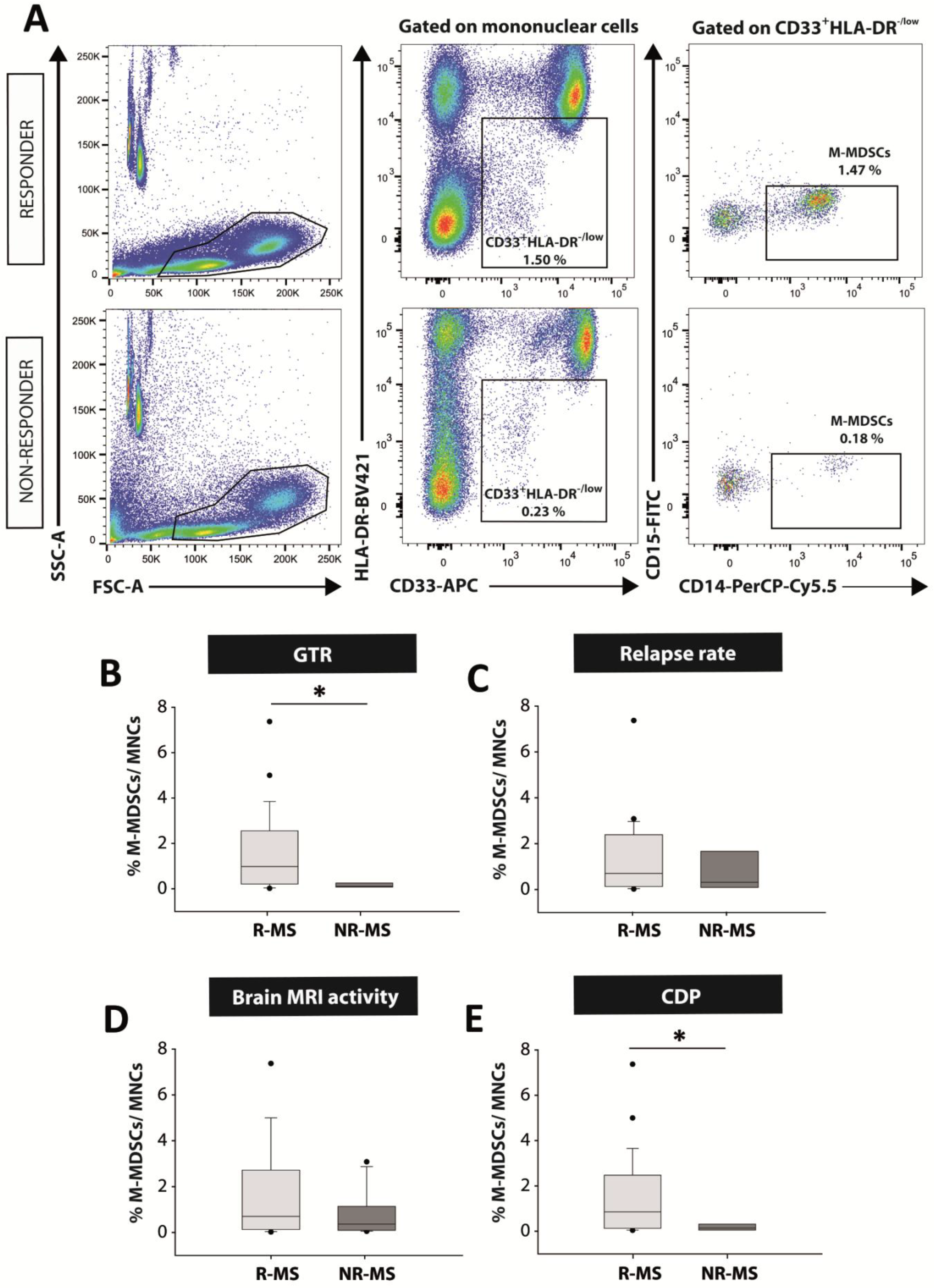
M-MDSCs are more abundant at baseline in patients with a good therapeutic response to fingolimod. **A**: Flow cytometry strategy of analysis for M-MDSCs in human PBMC samples. In both “responder” and “non-responder” MS patients, immature myeloid cells were gated from mononuclear subpopulation as CD33^+^HLA-DR^−/low^ populations. CD33^+^HLA-DR^−/low^ cells were assessed for CD14 and CD15 expression to identify the M-MDSC subset defined as CD14^+^CD15^-^. **B:** M-MDSCs were more abundant in R-MS patients (those who meet two or more criteria for NEDA-3: good therapeutic response –GTR) than NR-MS patients. **C-D**: M-MDSCs were similar between R-MS and NR-MS patients according the absence of new relapses (**C**), or new T2 lesions or Gd^+^ enhancing lesions in MRI after 12 months of treatment (**D**). **E**: M-MDSCs were clearly higher in patients with stable disability 12 months after fingolimod treatment. R-MS: “Responder”-MS patients; NR-MS: “non-responder”-MS patients. **B:** R-MS N =25, NR-MS N =6; in **C:** R-MS N =23, NR-MS N =8; in **D:** R-MS N =19, NR-MS N =12; in **E:** R-MS N =26, NR-MS N =5.

Finally, we assessed whether the level of M-MDSCs differed between R-MS and NR-MS for each of the clinical criteria used in the definition of NEDA-3. M-MDSCs were unable to distinguish R-MS and NR-MS patients based on the absence of new relapses or new MRI activity: *relapses*-R-MS 1.26 ± 0.35, NR-MS 1.14 ± 0.60 (p= 0.701); *MRI activity*-R-MS 1.50 ± 0.45, NR-MS 0.80 ± 0.29 (p= 0.441: Figure 7C-D). Importantly, the level of M-MDSCs was higher in MS patients with no evidence of an increase in disability (R-MS 1.43 ± 0.34, NR-MS 0.17 ± 0.07; p<0.05: Figure 7E).

In summary, these data displayed a clear parallel with the preclinical results reported previously in the EAE mice and they showed the ability of M-MDSC abundance to discriminate a good response to fingolimod treatment in MS patients. Moreover, M-MDSCs were enriched in patients whose disability remained stable or diminished after 12 months of fingolimod treatment.

## 3. Discussion

The present study demonstrates the potential of circulating Ly-6C^hi^ cells/M-MDSCs as a biomarker of response to fingolimod in both EAE mice and MS patients, respectively. From a preclinical point of view, our results show that peripheral Ly-6C^hi^ cell content at disease onset is the only immunological variable that allows us to distinguish the presence of a highly effective therapeutic response to fingolimod in the EAE model. M-MDSCs are also strongly represented in R-MS patients, irrespective of their demographic and baseline parameters. Furthermore, we identified that more circulating M-MDSCs are present at baseline in R-MS patients with no signs of CDP after a 12 month follow up.

The growing list of DMTs available for the treatment of MS [49] makes the identification of predictive biomarkers for treatment response ever more necessary to help clinicians adopt optimal treatment management strategies for MS patients [5]. In this context, promising results have been obtained from preclinical experimental procedures, although the translation of these findings to the clinic is often disappointing [50, 51]. This could be due to divergences between EAE and MS, as well as the more simplified interpretation of preclinical observations [52, 53]. To facilitate the translation of new potential biomarkers discovered in a preclinical setting into the clinical environment, the approximation of the EAE treatment strategy to the clinical reality of MS should be addressed. A large number of studies aimed at evaluating the efficacy of new treatments in EAE administer the drugs to animals prophylactically [54–56]. Conversely, a large amount of preclinical therapeutic data was obtained by treating at the same time point all EAE animals once they had reached a certain clinical score, or when only 50-60% of the animals had developed clinical signs of EAE [57–59], ignoring the inter-individual biological variability in disease severity found among immunized mice [24]. This type of approach is easier to follow as all animals are treated uniformly, monitoring is unified and the experimental procedure is completed simultaneously. However, these two experimental approximations are far from close to real clinical practice. In order to more closely resemble the clinical situation, we performed individualized administration and clinical assessment of each EAE mouse, and thus, vehicle or fingolimod treatment commenced at the time that each mouse began to develop clinical signs of EAE. Our data regarding the amelioration of the clinical course, the decrease in T cells and the histopathological preservation after fingolimod treatment demonstrated that this kind of experimental approach is as therapeutically valid as other preclinical designs [60, 61]. However, it also adds some valuable information about the inter-individual heterogeneity regarding the efficacy of the molecule tested, something closer to the reality of patient management.

Our data indicate that the level of Ly-6C^hi^ cells is a good and specific biomarker of clinical efficacy and responsiveness to fingolimod in EAE mice, i.e. it is useful to distinguish responder mice, and those that will have a less severe clinical development and fewer sequelae, following fingolimod treatment. The presence of experimental bias resulting from the randomized distribution of the groups can be ruled out since the timings, the level of clinical affectation and the immunoregulatory balance between M-MDSCs/T cells were similar between both experimental groups prior to treatment. Importantly, our case study is one of the few to demonstrate that EAE is very useful to test the responsiveness and efficacy of a given DMT. Most drugs that are successful in EAE fail to translate adequately into the clinic. This is due, amongst other reasons, from not taking into account the bimodal distribution of behavioral responses to drugs [62]. The nomenclature of “responders” and “non-responders” has been commonly used in MS patients for some time [14,63,64]. However, it is anecdotic in the study of new treatments in the EAE model [65, 66]. Our individualized study represents another exception since it mirrors the natural variability that exists among animals in response to fingolimod treatment, similar to that witnessed in MS patients. Data from this kind of experimental design will help identify so-called “disease classifying biomarkers”, which distinguish the MS endophenotypes that can yield measurable predictions in terms of prognosis and response to a corresponding treatment, a phenomenon that is crucial to improve future MS trials [67].

To our knowledge, this is the first study that describes the potential of Ly-6C^hi^ cells as biomarkers of response to DMTs in EAE. These findings are in agreement with our earlier observations demonstrating a clear relationship between the abundance of splenic Ly-6C^hi^ cells and the severity of the clinical course of EAE mice [23, 24]. In addition, it was reported that clinical improvements induced in EAE or other experimental models of MS through different experimental strategies increased the number and activity of splenic and CNS infiltrated Ly-6C^hi^ cells [32,43,68–71]. The seemingly controversial inverse relationship between the abundance of Ly-6C^hi^ cells (i.e. inflammatory monocytes/M-MDSCs) and clinical improvement in EAE has been explained by recent experimental observations. It is currently not possible to discriminate circulating Ly-6C^hi^ inflammatory monocytes from M-MDSCs in mice [18,20,21,41,42]. The vast majority of studies explored Ly-6C^hi^ function around the peak of EAE disease, when the greatest tissue damage has already occurred and Ly-6C^hi^ cells had acquired immunosuppressive functions, leading to a decrease in T cell proliferation and Th1/Th17 differentiation [41]. However, the deleterious role attributed to Ly-6C^hi^ cells in EAE results from their study during the pro-inflammatory phase of this animal model. In fact, an increase in these cells leads to an earlier onset and a worse disease severity [72], whereas their elimination resulted in milder disease courses [73].

To interpret these studies, it is important to consider the true functional consequences of Ly-6C^hi^ cells as they switch from behaving as antigen presenting cells at EAE onset to becoming T cell suppressors at the peak of the EAE clinical course [41]. Importantly, some studies identified peripheral blood Ly-6C^hi^ cells as precursors of M-MDSCs in certain tissues. At the peak of EAE, about 50% of the anti-inflammatory Arg-I^+^ monocyte-derived myeloid cells in the CNS are derived from iNOS^+^ pro-inflammatory cells that colonize this organ at disease onset [74]. Moreover, almost 98% of the Arg-I^+^ anti-inflammatory phagocytes infiltrating the CNS at the peak of the clinical course are thought to be derived from Ly-6C^hi^CCR2^+^ monocytes that invade the inflamed tissue from the onset of the disease [75]. Interestingly, it was shown that all the Arg-I^+^ cells present in the spinal cord at the peak of EAE have a phenotype of M-MDSCs and suppressor activity [76]. Along similar lines, the crucial role of pro-inflammatory Ly-6C^hi^ cells in effective repair at later stages of chronic inflammatory pathologies was reported recently [77]. All these data suggest that the vast majority of immunoregulatory phagocytic cells present in the CNS at the peak of EAE would be derived from circulating inflammatory monocytes that share their phenotype with M-MDSCs. Therefore, circulating Ly-6C^hi^ monocytes appear to be the only appropriate cell subset to be analyzed in order to validate M-MDSCs as a predictive tool for the response to fingolimod in EAE.

The promising preclinical results mentioned above and the need to find biomarkers that predict the response of the plethora of medications already approved to treat MS, would make M-MDSCs promising candidates to be used in clinical studies. To our knowledge, this is the first work that interrogates the role of M-MDSCs as predictive biomarkers of fingolimod treatment in MS. The only work that has explored the impact of DMTs on M-MDSCs indicated a non-significant increase in the number of total M-MDSCs in RR-MS patients treated with GA [32]. In fact, the scarcity of data about M-MDSCs and MS are somehow contradictory, and encourages a more in depth analysis on larger patient cohorts [31, 32]. To date, the analyses of immune cells as predictive biomarkers of the response to fingolimod centered on cells involved in adaptive immunity [14, 15],a s described for Ntz [78] or IFN-β [79]. By contrast, differences in cells of the innate immune response between those MS patients who respond efficiently to fingolimod or not has been restricted to regulatory CD56^bright^ NK cells, with a higher abundance in R-MS patients than in NR-MS patients [17]. However, the implication of myeloid cells as biomarkers of response to fingolimod has yet to be analyzed. In this sense, this study shows for the first time that the abundance of M-MDSCs at baseline is higher in MS patients that respond well to fingolimod. Moreover, the abundance of M-MDSCs in R-MS patients is mainly associated to the absence of CDP. Our results are consistent with our previous data regarding the higher abundance of circulating M-MDSCs in RR-MS patients at relapse relative to the lower EDSS at baseline and after a one year follow-up [80]. Therefore, EDSS seems to be the most relevant clinical parameter related to the enrichment of immunoregulatory M-MDSCs, the interpretation of which remains unclear and requires further experimental study.

In summary, our data strongly suggest that the percentage of circulating M-MDSCs seems to be a good parameter to identify patients with a higher probability of responding suboptimally to fingolimod treatment. This observation paves the way for the existence of good biomarkers for monitoring or predicting effective response to this or other DMTs in the same family (Ponsesimod, Siponimod, Ozanimod), which will facilitate their more accurately informed recommendation for patient management in the near future.

## Acknowledgements

The authors would like to thank Dr Virginia Vila-del Sol and Ángela Marquina Rodríguez at the Flow Cytometry Core Facility of the *Hospital Nacional de Parapléjicos* for their assistance with the phenotypic analysis of the different cell populations, and Dr José Ángel Rodríguez-Alfaro and Dr Javier Mazarío at the Microscopy Core Facility of the *Hospital Nacional de Parapléjicos* for their assistance with the confocal imaging and histological quantifications. Fingolimod was kindly supplied by Novartis Pharma AG under a signed Material Transfer Agreement.

## Author contributions

CC-T was a major contributor in writing the manuscript and carried out most of the experimental procedures. IM-D and JG-A contributed to the processing and analysis of the blood samples, both from the EAE mouse model and fingolimod treated patients. LC contributed to the processing and analysis of the PBMCs isolated from fingolimod-treated patients. MC performed the histopathological analysis in mice. DO, TC-T, LMV, LC-F, MC and LM isolated baseline (prior to the treatment) PBMCs samples from their own fingolimod patients, and performed the clinical evaluations. MCO contributed to the preclinical experimental design and contributed to the editing of the manuscript. JMG-D contributed to the clinical experimental design. DC designed the experiments, contributed to the data evaluation and analysis, and was a major contributor in writing and editing the manuscript. All authors read and approved the final version of the manuscript submitted.

## Data and material availability

The authors declare that all other data supporting the methods and the findings of this study are available within the paper as supplementary material.

