## Supplementary Tables for "Peripheral Myeloid-Derived Suppressor Cells are good biomarkers of the efficacy of Fingolimod in Multiple Sclerosis"

**Supplementary Table 1.** List of antibodies used for flow cytometry and histology

| Use | | Antibody | Target | Tissue/cells | Dilution[a] | Class | Clone | Manufacturer | Antibody ID |
|---|---|---|---|---|---|---|---|---|---|
| Flow cytometry | Mouse | CD11b-PerCP Cy5.5 | Myeloid cells | Peripheral Blood/ Splenocytes | 0.2μg | Rat monoclonal | M1/70 | BD Biosciences | AB_394002 |
| | | CD11c-APC | Dendritic cells | | 0.2μg | Hamster monoclonal | N418 | eBioscience | AB_469346 |
| | | F4/80-eFluor450 | Macrophages | | 0.2μg | Rat monoclonal | BM8 | eBioscience | AB_1548747 |
| | | MHC-II-PE-Cy7.7 | Antigen presenting cells | | 0.2μg | Rat monoclonal | M5/114.15.2 | eBioscience | AB_10870792 |
| | | Ly-6C-FITC | MDSCs | | 0.2μg | Rat monoclonal | Ly-6C:AL-21 | BD Biosciences | AB_394628 |
| | | Ly-6G-PE$^-$ | | | 0.2μg | Rat monoclonal | Ly-6C:1A8 | BD Biosciences | AB_394208 |
| | | CD3-PB | T cells | | 0.2μg | Hamster monoclonal | 500A2 | BD Biosciences | AB_397063 |
| | | CD4-PE | CD4$^+$-T cells | | 0.1μg | Rat monoclonal | RM4-5 | BD Biosciences | AB_394585 |
| | | CD8-FITC | CD8$^+$-T cells | | 0.25μg | Rat monoclonal | 53-6.7 | BD Biosciences | AB_394568 |
| | | CD25-PE-Cy5.5 | Early activated T cells | | 0.2μg | Rat monoclonal | PC61.5 | eBioscience | AB_11218898 |
| | | CD69-APC | Early activated T cells | | 0.2μg | Hamster monoclonal | H1.2F3 | eBioscience | AB_1210795 |
| | Human | CD11b-PE-Cy7 | MDSCs | Peripheral Blood | 0.25μg | Mouse monoclonal | ICRF44 | BD Biosciences | AB_396849 |
| | | CD33-APC | | | 0.2μg | | WM53 | BD Biosciences | AB_398502 |
| | | HLA-DR-BV421 | | | 0.2μg | | BM8 | BD Biosciences | AB_2687421 |
| | | CD14-PerCP-Cy5.5 | | | 0.25μg | | MΦPg | BD Biosciences | AB_2737726 |
| | | CD15-FITC | | | 0.2μg | | HI98 | BD Biosciences | AB_395801 |
| Histology | Human/ mouse | NF-H | Non-phosphorylated neurofilament protein | Spinal cord | 1:200 | Mouse monoclonal | SMI-32 | Biolegend | AB_2564642 |

[a]Refers to the amount per million cells. Abbreviations: FITC, fluorescein isothiocyanate; PB, pacific blue; PE, phycoerythrin.

**Supplementary Table 2.** Demographic data from the fingolimod patients included in the study.

| | Total cohort (N = 31) | R-MS (N = 25) | NR-MS (N = 6) | Ntz patients (N = 11) | Non-Ntz patients (N = 20) |
|---|---|---|---|---|---|
| ***Age (years)*** ¥ | 39.5 ± 1.31 | 40.3 ± 1.37 | 36.3 ± 3.67 | 38.8 ± 1.61 | 41.0 ± 2.31 |
| ***Sex (% of female)*** $ | 26 (83.87%) | 21 (84%) | 5 (83.33%) | 17 (85%) | 9 (81.82%) |
| ***Previous DMTs*** | 1 naïve<br>15 IFNb<br>(5 IFNb-1b, 10 IFNb-1a)<br>4 GA<br>11 Ntz | 1 naïve<br>14 IFNb<br>(4 IFNb-1b, 10 IFNb-1a)<br>3 GA<br>7 Ntz | 1 IFNb-1b<br>5 Ntz | - | 1 naïve<br>15 IFNb<br>(5 IFNb-1b, 10 IFNb-1a)<br>4 GA |
| ***0 treatment*** | 1 | 1 | 0 | 0 | 1 |
| ***1 treatment*** | 12 | 9 | 3 | 2 | 10 |
| ***2 treatment*** | 4 | 4 | 0 | 0 | 4 |
| ***≥3 treatment*** | 14 | 11 | 3 | 9 | 5 |
| ***Basal ARR*** ¥,˜ | 1.09 ± 0.18 | 1.24 ± 0.19 | 0.50 ± 0.34 | 0.27 ± 0.14 | 1.55 ± 0.20 * |
| ***Basal EDSS*** // | 3.0 (2.0-4.0) | 3.0 (2.0-4.13) | 3.0 (2.5-4.0) | 3.0 (2.13-4.0) | 3.0 (2.0-4.25) |
| ***Gd+ lesions*** ¥ | 1.45 ± 0.54 | 1.64 ±0.66 | 0.67 ± 0.422 | 2.15 ± 0.80 | 0.18 ± 0.18 * |

*Abbreviations: ARR, Annualized Relapse Rate; GA, Glatiramer acetate; IFN, Interferon; NR-MS, Non-responders MS patients; Ntz, Natalizumab; R-MS, Responder MS patients;*

*¥ The values are the mean ± SEM of each group.*

*// The values are the median IQR.*

*˜ For basal ARR, only the last year was considered.*

** Significant differences after Mann-Whitney test versus Ntz.*

*$ A Fisher exact test was used to compare the proportions.*

**Supplementary Table 3**. Correlations between M-MDSCs and clinical parameters after 12 months of fingolimod treatment.

| | EDSS | ΔEDSS | New T2 lesions | New $Gd^+$ lesions | Number of Relapses | ΔRelapses |
|---|---|---|---|---|---|---|
| **Total cohort** (N =31) | r = -0.085<br>p = 0.648 | r = -0.201<br>p = 0.276 | r = -0.137<br>p = 0.459 | r = -0.081<br>p = 0.664 | r = -0.115<br>p = 0.534 | r = -0.262<br>p = 0.153 |
| **R-MS$_{NEDA-3}$** (N = 13) | r = 0.149<br>p = 0.616 | r = 0.190<br>p = 0.516 | - | - | - | r = 0.129<br>p = 0.669 |
| **R-MS$_{CR}$** (N = 20) | r = 0.108<br>p = 0.644 | r = 0.098<br>p = 0.676 | r = 0.151<br>p = 0.517 | r = 0.107<br>p = 0.649 | - | r = -0.092<br>p = 0.695 |
| **R-MS** (N = 25) | r = 0.043<br>p = 0.835 | r = 0.091<br>p = 0.660 | r = 0.107<br>p = 0.607 | r = 0.095<br>p = 0.649 | r = 0.167<br>p = 0.423 | r = -0.010<br>p = 0.960 |

*A Spearman's correlation was carried out.*
